# Towards best-practice approaches for CRISPR/Cas9 gene engineering

**DOI:** 10.1101/469544

**Authors:** Claude Van Campenhout, Pauline Cabochette, Anne-Clémence Veillard, Miklos Laczik, Agnieszka Zelisko-Schmidt, Céline Sabatel, Maxime Dhainaut, Benoit Vanhollebeke, Cyril Gueydan, Véronique Kruys

## Abstract

In recent years, CRISPR has evolved from “the curious sequence of unknown biological function” into a functional genome editing tool. The CRISPR/Cas9 technology is now delivering novel genetic models for fundamental research, drug screening, therapy development, rapid diagnostics and transcriptional modulation. Despite the apparent simplicity of the CRISPR/Cas9 system, the outcome of a genome editing experiment can be substantially impacted by technical parameters as well as biological considerations. Here, we present guidelines and tools to optimize CRISPR/Cas9 genome targeting efficiency and specificity. The nature of the target locus, the design of the single guide RNA and the choice of the delivery method should all be carefully considered prior to a genome editing experiment. Different methods can also be used to detect off-target cleavages and decrease the risk of unwanted mutations. Together, these optimized tools and proper controls are essential to the assessment of CRISPR/Cas9 genome editing experiments.

## Introduction

Engineered nucleases, from zinc-finger nucleases to TALENs and CRISPRs, form a powerful class of genome editing tools. Among these, the CRISPR/Cas9 system has become the most popular, owing to its ease of use and rapidity. The CRISPR/Cas system was discovered in prokaryotes where it provides adaptive immunity against foreign elements (Mojica et al., 2005; Barrangou et al., 2007; Deltcheva et al., 2011; Gasiunas et al., 2012). In 2013, the CRISPR/Cas9 system from *Streptococcus Pyogenes* (spCas9, further indicated in the text as Cas9) was successfully adapted for genome editing in eukaryotic cells (Cong et al., 2013, Cho et al. 2013; Jinek et al. 2013; Mali et al., 2013). Since then, the technique has become extremely popular as it can modify the genome of a large variety of organisms with unprecedented ease.

Despite the potential of the CRISPR technology, not all genome editing experiments work equally well and this technology is not as easy as it was once assumed. Despite significant improvements, there is still limited predictability of whether the CRISPR system will be able to effectively target a given region of interest. This aspect is of particular importance in the context of CRISPR/cas9-based screens in model organisms and is related to the definition of the target site and the sequence of small guide RNA (sgRNA). Another major hurdle common to all engineered nucleases is the risk of unwanted mutations at sites other than the intended on-target site (off-target effects). The off-target mutations are the consequence of sgRNA binding to DNA sites with less than perfect complementary (Fu et al., 2013; Hsu et al., 2013). Current strategies to increase targeting specificity notably include: refinements in guide RNA selection, enzyme and guide engineering and improvements in the delivery method. Here, we describe a series of guidelines to optimize CRISPR/Cas9 efficiency and specificity.

### Analysis of the target locus

Careful determination of target sites is essential. For many applications, a loss of function may be desirable or even required. Targeting of functional protein domains was recently shown to result in higher proportions of loss-of-function mutations (Shi et al., 2015). A common strategy is to select sgRNAs that will target Cas9 nuclease to the N-terminal-coding exons of protein coding genes. After the action of Cas9 nuclease, the introduction of indels by the error-prone Non-Homologous End Joining repair of double strand breaks introduces frame-shift mutations and subsequent premature stop codons, leading to mRNA elimination by nonsense-mediated mRNA decay. Genome editing experiments to generate knockouts should be designed to disrupt exons that are shared by all transcript variants of a given gene. This strategy can also be applied to whole gene families using a sgRNA against exons that are conserved between all family members (Endo M et al., 2015). The CRISPys algorithm aims at designing the optimal sequence to target multiple members of gene family (Hyams et al., 2018).

The high frequency with which CRISPR-induced mutations can be directed to target genes enables easy isolation of homozygous gene knockouts. Paradoxically, a potential caveat is found in this high efficiency. This holds particularly true in cell lines upon targeting genes essential for cell viability and fitness. In this regard, two distinct genome-wide CRISPR-Cas9-based screens have identified ≈2000 essential genes in the human genome (Hart et al., 2015; Wang T et al., 2015). More recently, Lenoir and colleagues published a database of pooled *in-vitro* CRISPR knockout library essentiality screens that can be searched to identify genes which are essential across different human tissues (Lenoir et al., 2018).

Genetic screen in zebrafish and mouse have estimated that as many as 30% of genes are embryonic lethal (Driever et al. 1996; Haffter et al., 1996; Ayadi et al., 2012). The functional characterization of such essential genes requires the generation of heterozygous knockouts. The generation of hypomorphic alleles with the CRISPR system has been reported by different groups (Challa et al., 2016; Goto et al., 2016) but the method is not, at the moment, commonly used.

RNAi or CRISPRi (Qi et al., 2013) are efficient alternative loss-of-function methods and their effects can be directly evaluated at the transcriptome level. In addition, the development of inducible CRISPR tools provides a solution for genome editing with tight temporal control (Zestche et al., 2015b; Cao et al., 2016). They additionally circumvent the mechanisms of genetic compensation that not unfrequently mask the phenotypes of knock-outs but not knockdown models (El-Brolosy and Stainier, 2017).

Genetic polymorphism in the target region should be carefully assessed as it might have a profound influence on the CRISPR/Cas9 efficacy. Although base mismatches (up to 5) may be tolerated between the sgRNA and targeted sequences, the PAM and its proximal sequence have a stricter adherence to the consensus (Zheng et al., 2017). When a sgRNA is selected, the potential presence of a single-nucleotide polymorphism (SNP) in the PAM and the sgRNA-binding site should be verified as it can abolish Cas9 binding and cleavage. Of note, commonly used laboratory cell lines such as HeLa present a variant spectrum that slightly differs from the one found in the human population (Landry et al., 2013). In general, sequences found in genomic databases may not exactly correspond to the DNA sequences of the model used for the genome editing experiment. Sequencing of the target locus prior to sgRNA design will solve this potential pitfall. On the opposite, this PAM constraint can be exploited to target and disrupt heterozygous single-nucleotide mutations in certain dominant autosomal disorders, while leaving the wild-type allele intact (Courtney et al., 2016; Li et al., 2016). Cell line ploidy is an additional consideration to take into account. Many common laboratory cancer cell lines carry four or more copies of a chromosome. Full knockouts would then require the introduction of mutations in all copies of the target gene. In practice, it is strongly advised to sequence the target loci to verify homozygous knockout when generating mutant clonal cell lines (see further section).

Besides the influence of the sequence features, chromatin states also strongly impact Cas9 binding and nuclease activity in vertebrates. Nucleosomes constitute fundamental units of chromatin and their positioning directly impedes Cas9 binding and cleavage *in vitro* and *in vivo*. Highly active sgRNAs for Cas9 are found almost exclusively in regions of low nucleosome occupancy (Horlbeck et al., 2016; Isaac et al., 2016). Higher order chromatin structure (i.e. organization beyond the level of the linear array of nucleosomes) also influences Cas9 binding and enzymatic activity. Several authors showed that Cas9 cleavage efficiency positively correlates with open chromatin based on DNase I hypersensitivity. Along the same line, the activity of Cas9 can be significantly hindered by compact heterochromatin in cells (Daer et al., 2017). Interestingly, the engineered Cas9 variants designed to improve specificity, Cas9-HF1 (Kleinstiver et al., 2016) and eSpCas9(1.1) (Slaymaker et al., 2016), might be even more impacted than Cas9 by the chromatin-related factors (Chen et al., 2016; Jensen et al., 2017; Chen et al., 2017). While some gene editing applications have the option to select easy-to cleave targets, such practice may not be feasible for gene corrections and other potential therapeutic applications. Many CRISPR genome editing experiments focus on gene targeting and the study of the phenotypic consequences. In these applications, the gene of interest is usually transcriptionally active and the associated chromatin is relatively accessible to Cas9. Nevertheless, chromatin compactness can vary considerably between different genomic sites and from one cell type to another. Gene targeting in model organisms presents an additional challenge as chromatin landscape is under constant change to ensure coordinated growth and differentiation during early development. Atlases of transcriptional activity (RNA-Seq) and of chromatin accessibility (ATAC-Seq, ChIP-Seq, …) are valuable resources of information (see notably the ENCODE project: https://www.encodeproject.org) to predict sgRNA efficiency (Uusi-Makela et al., 2018) and have been used to elaborate a predictive algorithm for zebrafish sgRNA selection taking into account chromatin accessibility (Chen et al., 2017). Gene editing in mouse and human cells has been greatly facilitated by the publication of the genome-wide Brie and Brunello libraries (Doench et al., 2016). These optimized sgRNA libraries respectively target the mouse and human genomes, and provide 3-4 sgRNA sequences per gene with predicted high on-target efficiency and low off-target effects.

For more challenging applications such as the editing of heterochromatin embedded sequences, chromatin manipulation might enhance the CRISPR targeting efficiency. While treatment with chromatin-disrupting drugs does not appear sufficient, transient overexpression of a targeted transcriptional activator might be an effective method to enhance Cas9 editing at closed chromatin regions (Daer et al., 2017).

### Delivery methods

Introduction of the CRISPR/Cas9 components into cultured cells is often achieved by DNA-based delivery systems such as transfection of plasmids encoding nuclear targeted Cas9 and sgRNA. Transduction with viral particles is also commonly used and is typically more efficient compared to plasmid transfection and is applicable to many cell types including primary cells. Plasmid transfection and viral transduction methods lead to a prolonged or a permanent expression of Cas9, respectively. Extended expression of Cas9 in cells can lead to accumulation of off-targeting events (Kim et al., 2014). Indeed, constitutive expression of lentiviral-based Cas9 and sgRNAs leads to an enrichment of predicted off-target sites over time. Reducing the concentration of delivered plasmid during transfection was shown to decrease off-targeting (Hsu et al., 2013). These data support the idea that controlling the expression of Cas9 and the sgRNAs in order to limit the time of action can reduce genome-wide off-targeting. A doxycycline-inducible promoter allows for transient Cas9 expression and is compatible with lentiviral delivery of the nuclease (Wang et al., 2018). Because gene editing results in a permanent change in the genome, CRISPR-mediated editing can be achieved using Cas9 protein/sgRNA ribonucleoprotein (RNP) complexes. sgRNAs can be rapidly synthetized *in vitro* or ordered from various sources. Recombinant nuclear targeted Cas9 protein can also be produced in house or obtained from different commercial suppliers. RNP complexes can be delivered by a variety of techniques such as lipid-mediated transfection (Liang et al., 2015; Zuris et al., 2015), electroporation (Kim et al., 2014), induced transduction by osmocytosis and propanebetaine (iTOP) (D’Astolfo et al., 2015), micro-injection (Gagnon et al., 2014) or cell-penetrating peptide-mediated delivery (Ramakrishna et al., 2014a). Uncoupling administration of the sgRNA and Cas9 protein (e.g. in the context of genome-scale screens) can lead to successful gene editing in human primary cells (Shifrut et al., 2018) and appears to be more efficient upon delivery of Cas9 protein complexed with a non-targeting gRNA (Ting et al., 2018). Finally, biolistic transfer of Cas9/sgRNA RNP complexes or of Cas9- and sgRNA-encoding plasmids appears as an attractive alternative for cells resistant to other delivery methods such as plant cells (Svitashev et al., 2016; Hamada et al., 2018).

The sgRNA-Cas9 RNPs were shown to cleave the target chromosomal DNA between 12 and 24 h after delivery and the frequency of gene editing reaches a plateau after one day. For plasmid expression of Cas9 and sgRNA, equivalent gene editing levels were only achieved at three days after delivery (Kim et al., 2014). Furthermore, the Cas9 protein has been shown to be degraded rapidly in cells, within 24–48 h after delivery, compared to several days when continuously expressed from a plasmid (Kim et al., 2014; Liang et al., 2015). Several authors showed that the ratio of the indel frequency at the on-target site to off-target sites strongly increases when RNPs are transfected in comparison with plasmid (Ramakrishna et al., 2014a; Liang et al., 2015). While off-target effects may be less of a concern in screening applications since any identified “hits” will be confirmed through follow-up experiments, constitutive expression or high stability of Cas9 nuclease and/or sgRNA may be undesirable for many applications, such as generation of clonal cell lines for a phenotypical study of a specific gene knockout. In addition to the increased potential for off-target effects due to prolonged or constitutive expression of components of the CRISPR-Cas9 system, unwanted incorporation of the plasmid DNA into the cell genome is not uncommon. When the DNA repair pathways are activated after Cas9-mediated double-stand breaks, the risk of foreign DNA integration is increased. The absence of transgene eliminates the risk of unintended DNA integration.

The delivery of RNP complexes has also the major advantage to be easily applicable to a wide range of model organisms and cell types. In *in vivo* contexts, the functionality of the Cas9-sgRNA RNP complexes has been reported as being superior to other delivery methods. In zebrafish, mutagenesis can be performed through micro-injection of Cas9-encoding mRNA or of Cas9 protein together with sgRNA into fertilized embryos. Contrary to Cas9-encoding mRNA, RNPs are immediately active upon microinjection and are generally more effective (Burger et al., 2016 and **Figure 1**). This is of significant importance as the first cell division in zebrafish occurs very rapidly (40 min. after fertilization) and mutagenesis occurring after this first division leads more likely to mosaicism. The fact that sgRNAs can be easily synthetized *in vitro* makes it possible to use multiple sgRNAs simultaneously to achieve multigenic targeting (Liang et al., 2015; Song et al., 2017). The DNA-free system also suppresses the variability that can arise from the choice of promoter used to drive expression from vector-based CRISPR-Cas9 systems. It is well known that not all promoters are functional in every cell type or cell line, so delivery of Cas9 protein or Cas9 mRNA avoids incompatibilities of certain promoters in specific cells. Codon usage patterns also vary between species and Cas9 derived from DNA or mRNA expression may not yield the expected result as every organism has its own codon bias. Optimization of codon usage is a routine process but can be relatively time-consuming. Codon optimization becomes unnecessary when using Cas9 protein instead of a DNA- or mRNA-based delivery method. Independent of the delivery mode, specific anti-Cas9 antibodies can be used to measure Cas9 expression level by western and to confirm Cas9 presence in the nucleus (**Figure 2**).

**Figure 1.**
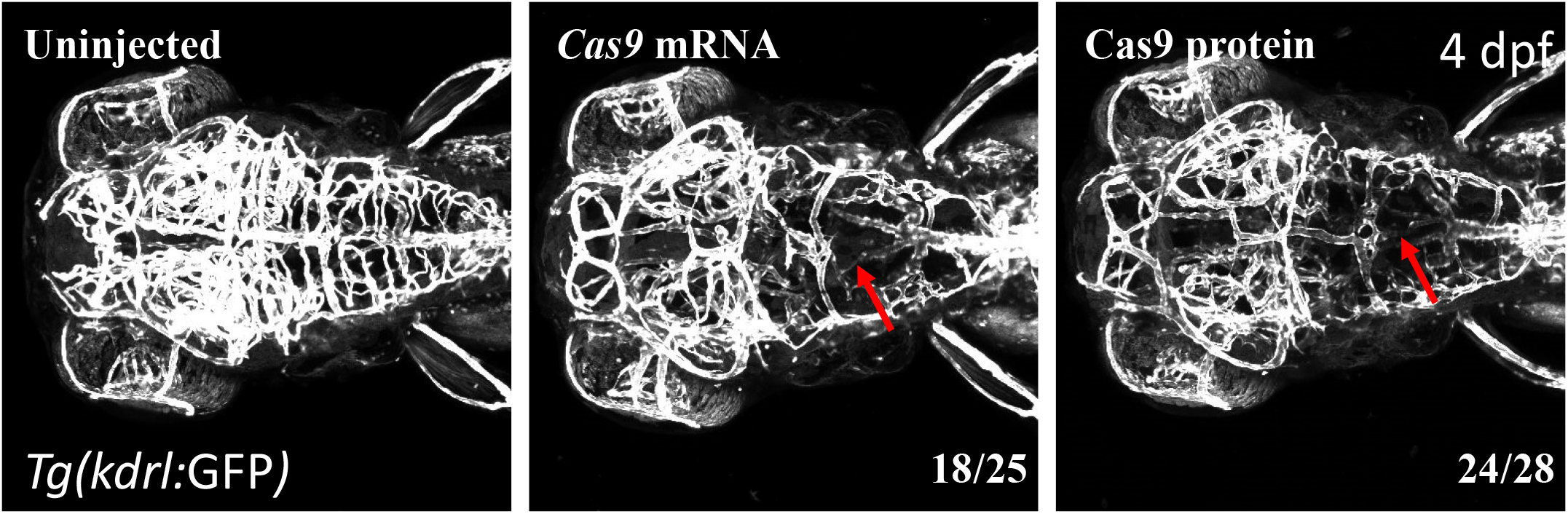
Cas9 Nuclease protein NLS and *Cas9* mRNA injections achieve high efficient bi-allelic somatic gene disruptions in zebrafish. *Tg(kdrl:GFP)^s843^* embryos were injected at the one-cell stage with 300 pg of recombinant Cas9 protein (Cas9 Nuclease Protein NLS from Diagenode, cat n° C29010001) or 150 pg of synthetic *cas9* mRNA obtained by *in vitro* transcription from the XbaI linearized pT3TS-nls-zcas9-nls vector, Addgene #46757 using the mMessage mMachine T3 kit (Ambion)) and 30 pg of two guide RNAs targeting *reck*, an essential regulator of central nervous system vascularization (Vanhollebeke et al., 2015; Ulrich et al., 2015; Eubelen et al., 2018). The two reck-targeting sgRNAs constructs were obtained by *in vitro* annealing of the following primers: 1-Fwd: 5’-TAGGCCTGACAGTACTCACGAC-3’; 1-Rev: 5’-AAACGTCGTGAGTACTGTCAGG-3’; 2-Fwd: 5’-TAGGTGCGCAGGATACGCTTAC-3’; 2-Rev: 5’-AAACGTAAGCGTATCCTGCGCA-3’ and cloning into pT7-sgRNA (Addgene #46759) as described in Jao et al. 2013. sgRNAs were transcribed from the BamHI linearized vector pT7-sgRNA using the MEGAshortscript T7 kit (Thermo Fisher Scientific). Maximal intensity projections of confocal z-stacks of the cranial vasculature of *Tg(kdrl:GFP)* larvae at 4 days post-fertilization (dpf) Red arrows point to the avascular hindbrains. The fraction of larvae showing cerebrovascular defects is indicated in each panel.

**Figure 2.**
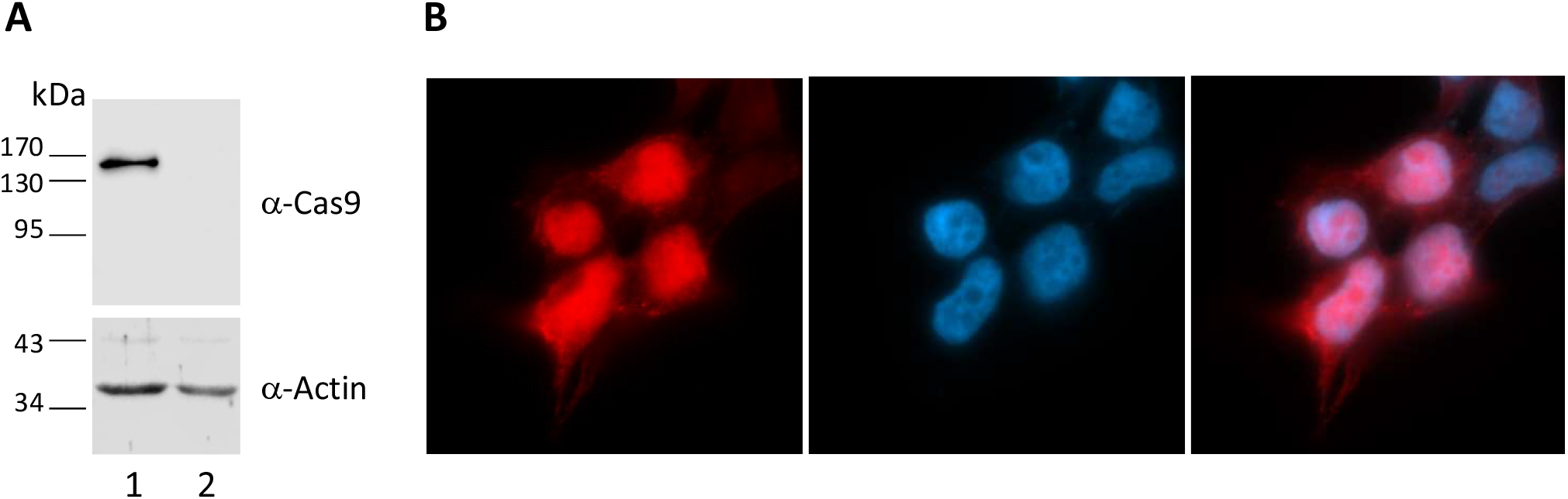
Cas9 expression can be detected by western blot and immunofluorescence. **A.** Western blot analysis of HEKT293T cells transfected with pSpCas9(BB)-2A-Puro (PX459) V2.0 plasmid (Addgene #62988, Ran et al., 2013) encoding Cas9 using Lipofectamine 2000 (Life Technologies). Cell lysates from transfected (lane 1) or untransfected cells (lane 2) were analyzed by immunoblotting with mouse anti-Cas9 (Diagenode Cat No. C15200216). Equal amounts (5 μg) of proteins were loaded in each lane and protein bands were visualized using anti-mouse HRP-coupled sheep IgG (GE Healthcare). Actin was analyzed as loading control. HRP-induced chemiluminescence was measured on a Licor Odyssey FC imaging system using supersignal West pico (Thermo Scientific) chemiluminescent substrate. **B.** HEK293T cells were transfected with dCas9 plasmid (Addgene #100091, O’Geen et al., 2017). dCas9 localization was analyzed by immunostaining using anti-Cas9 mouse monoclonal antibody (Diagenode Cat No. C15200216) and Alexa-Fluor 594-conjugated donkey anti-mouse antibody. Nuclei were counterstained with DAPI.

Lentiviral or plasmid delivery of Cas9 and sgRNA often utilizes a selection gene encoding either a drug selectable marker (hygromycin, blasticidin, puromycin, …) or a reporter protein (GFP, NGFR, …) to isolate cells that are successfully transduced or transfected. When RNPs are transfected or electroporated, alternative strategies can be used such as surrogate reporters (Ramakrishna et al., 2014b, He et al., 2016; Wu et al., 2017). However, these methods are inefficient for assessing sgRNAs efficiency at a large scale because it is both time- and labour-consuming to construct a specific reporter for each individual sgRNA. To avoid specific cloning, the transfection efficiency can also be indirectly evaluated with a dTomato reporter assay (D’Astolfo et al., 2015). Fluorescent versions of Cas9 such as Cas9-GFP (Mircetic et al., 2017) or Cas9-Cy3 (Kim et al., 2017) can also be used to sort RNPs-transfected cells. These latter methods focus on the physical separation of edited cells from unedited cells. An important aspect to consider is that CRISPR experiments lead to a genetic heterogeneity due to the random nature of DNA repair by the NHEJ pathway. As this genetic heterogeneity could yield phenotypic heterogeneity, monoclonal populations should be isolated prior to phenotypic analysis. The first step is to determine the editing efficiency of the entire cell population. This information can indicate how many individual clones should be isolated and checked for editing. If limited dilutions are used to isolate individual cells, it should be realised as soon as possible after termination of the edition process, as non-edited cells could potentially outgrow edited cells.

Gene editing in *in vivo* mouse models was greatly facilitated by the generation of a knock-in (KI) transgenic mouse in which a Cre-inducible Cas9-P2A-GFP was inserted in the Rosa26 locus (Platt et al., 2014). Cre-recombination leads to cell- or tissue-specific Cas9 expression, as evidenced by GFP expression. Apart from allowing for gene editing following *in vivo* delivery of sgRNAs, this model can also be used to efficiently edit the genome of primary cells *ex vivo*.

### Guide RNA efficiency and specificity

The performance of sgRNAs targeting the same gene can vary dramatically. This was recently highlighted in a novel approach to CRISPR genomics where expression of sgRNAs was coupled with specific protein barcodes, allowing for simultaneous multidimensional phenotypic analysis of several dozens of knockouts at a single cell resolution (Wroblewska et al, 2018). In a pooled parallel analysis of gene editing efficiency for 10 genes (3-4 sgRNAs guide per gene), the authors demonstrated that the gene KO at the protein level was highly variable depending on the sgRNAs used. There are many bioinformatic tools available for sgRNA design and some of these tools also apply filters or show ‘scores’ related to predicted effectiveness. Small guide RNAs with potential for weak secondary structures are likely to be more efficient than alternatives with strong secondary structures (Thyme et al., 2016). Nevertheless, no computational tool can guarantee the efficacy of a sgRNA and, when possible, several sgRNAs should be tested. Endonuclease cleavage assays can be used to characterize the *in vitro* efficacy of a particular sgRNA. Experimental validation of sgRNAs before practical application is particularly important to minimize wasted experiments on sgRNAs with poor activity. In these *in vitro* assays, the target DNA site, including its PAM motif, is either inserted into a plasmid or provided in the form of a PCR product. The Cas9 recombinant protein and the sgRNA are pre incubated in a 1:1 molar ratio in the cleavage buffer to reconstitute the Cas9-sgRNA complex prior to the addition of target DNA. Cleavage of plasmid or PCR substrates are monitored by agarose gel electrophoresis with an intercalating dye (**Figure 3**). The reaction rate can strongly vary in function of DNA source and length (PCR product versus plasmid, circular plasmid versus linear plasmid), optimal enzyme and substrate concentrations, and also reaction time points need to be determine empirically (Anders and Jinek, 2014). This *in vitro* test validates sgRNA intrinsic capacity to form cleavage-competent complexes, however it does not guarantee *in* vivo effectiveness which also greatly depends on chromatin accessibility as previously mentioned.

**Figure 3.**
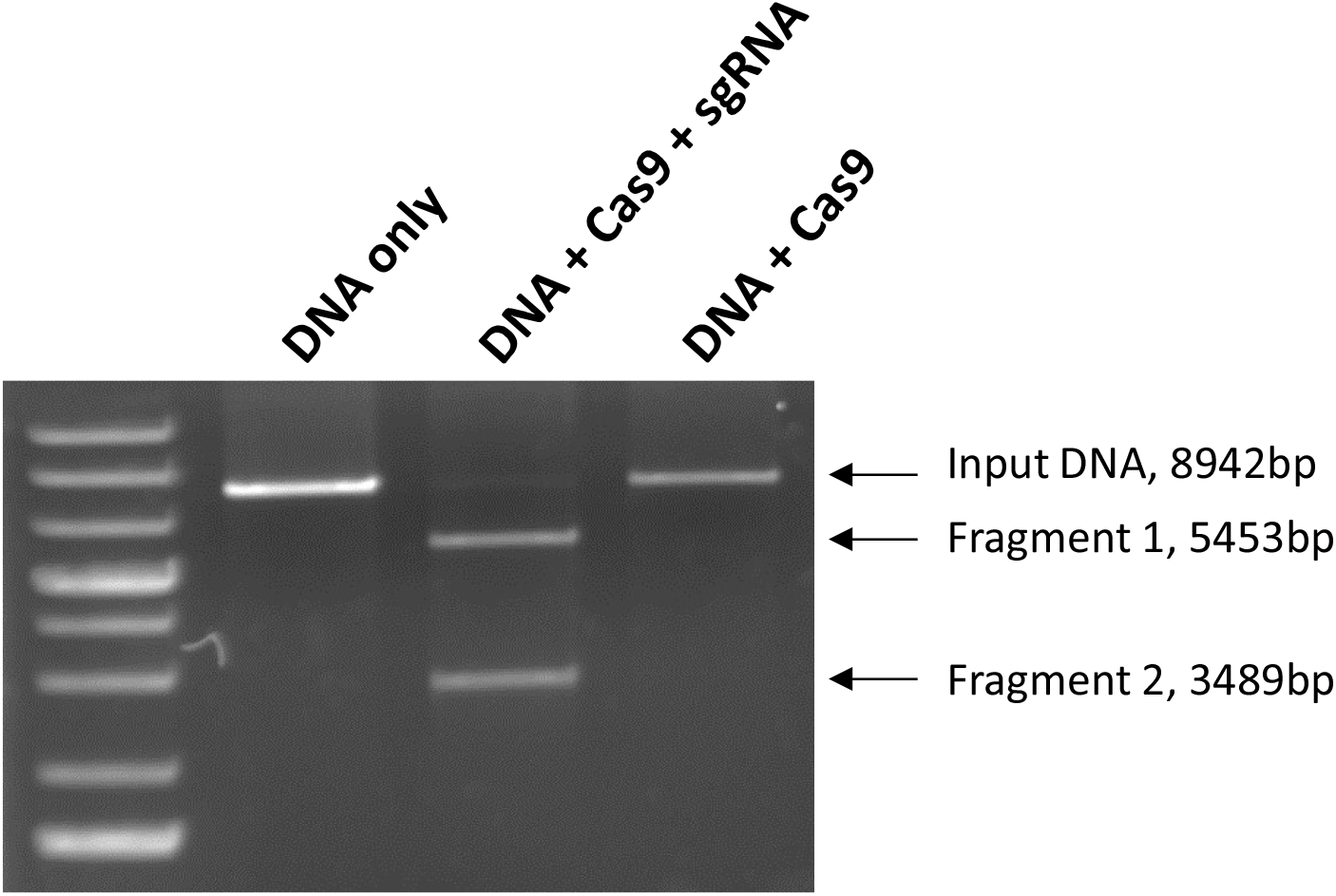
*In vitro* validation of sgRNA by Cas9 cleavage assay. DNA target is PvuI-linearized CRISPR-SP-Cas9 reporter plasmid (Addgene #62733, D’Astolfo et al., 2015). Corresponding sgRNA (GGGCCACUAGGGACAGGAU) was synthetized by *in vitro* transcription. Target DNA was incubated with (lanes 2 and 3) or without (lane 1) Cas9 recombinant protein (Cas9 Nuclease protein NLS from Diagenode, Cat No. C29010001). Reactions were set-up with a ratio of 20:20:1 (Cas9:sgRNA:DNA target) and incubated 1 hour at 37°C. The products were resolved on 0.8% TAE agarose gel stained with ethidium bromide.

The targeting specificity of Cas9 is believed to be tightly controlled by the 20-nt guide sequence of the sgRNA and the presence of a PAM adjacent to the target sequence in the genome. Nevertheless, potential off-target cleavage activity can still occur on DNA sequence with even three to five base pair mismatches in the PAM-distal part of the sgRNA-guiding sequence (Fu et al., 2013). Of note, shortening of sgRNA guide sequence to 17-18 nucleotides was shown to improve target specificity (Zhang et al., 2016; Fu et al., 2014). Numerous online tools are available to assist in sgRNA design but the correlation between the predictions and the actual measurements vary considerably since sequence homology alone is not fully predictive of off-target sites (Haeussler et al., 2016). These tools also suggest probable off-target sites but the appropriate number of potential sites to experimentally assay remains unclear. Moreover, there are still contradictory conclusions as to the prevalence of off-target effects, from low (Kim et al., 2015) to high levels of off-targeting (Tsai et al., 2015).

Cleavage at on- and off-target sites can be assessed using various methods which include mismatch-sensitive enzymes (Surveyor or T7 endonuclease I assay), restriction fragment length polymorphism (RFLP analysis), High Resolution Melting curve Analysis (HRMA) or PCR amplification of the locus of interest followed by sequencing. Surveyor and T7 Endonuclease I specifically cleave heteroduplex DNA mismatch. The T7 endonuclease I assay outperforms the Surveyor nuclease in terms of sensitivity with deletion substrates, whereas Surveyor is better for detecting single nucleotide changes. The limit of sensitivity for T7 endonuclease I assay is around 5% (Vouillot et al., 2015). HRMA utilizes the difference in *melting* curve of the heteroduplex and mutant homoduplex. A recent report demonstrates that techniques such as targeted Next Generation Sequencing (NGS), Tracking Indels by Decomposition (TIDE) and Indel Detection by Amplicon Analysis (IDAA) outperform nuclease-based methods to detect Cas9-mediated edition in pools of cells (Sentmanat et al., 2018). Ultimately, Sanger sequencing of DNA from individual clones is the gold standard for confirming the presence of indels at on-target site but is not easily applicable to off-target detection. Overall, these indels detection methods are relatively straightforward but are low throughput and interrogate one locus at the time.

Unbiased off-target analysis requires the detection of mutations generated in the target cells by the CRISPR/Cas9 system outside their target locus. In theory, Whole Genome Sequencing (WGS) of cells before and after editing could be used to study CRISPR/Cas9 specificity. In a clonal population, off-target sites can be determined by the analyses of the new mutations that have been generated outside the intended locus. However, WGS faces its own challenges and might not be easily applicable to the detection of off-target mutations. While sequencing costs continue to drop, a certain degree of bioinformatic expertise is necessary to detect small indels and separate signal from noise. In fact, many spontaneous new mutations may appear during clonal expansion and it might not be possible to distinguish them from off-target effects. WGS of individual induced Pluripotent Stem Cells clones reveals a large number of indels in the genome that are not the result of Cas9 activity, but rather a consequence of clonal variation or technical artefacts (Smith et al., 2014). To circumvent these limitations, several methods have been recently developed to measure Cas9 off-target activity across the genome such as BLESS (labeling of double strand breaks followed by enrichment and sequencing) (Ran et al., 2015), HTGTS (high throughput genome-wide translocation sequencing) (Frock et al., 2015), GUIDE-Seq (genome-wide unbiased identification of double-strand breaks enabled by sequencing) (Tsai et al., 2015), Digenome-Seq *(in vitro* Cas9-digested whole genome sequencing) (Kim et al., 2015), IDLV (detection of off-targets using integrase-deficient lentiviral vectors) (Wang X et al., 2015) and most recently, SITE-Seq (a biochemical method that identifies DNA cut sites) (Cameron et al., 2017) and CIRCLE-Seq (an *in vitro* method for identifying off-target mutations) (Tsai et al., 2017). Overall, these unbiased methods tend to be less sensitive and have a lower throughput than biased targeted sequencing, in addition to typically requiring higher sequencing coverage and much more complex protocols. These techniques also require manipulation of the genome and might be difficult to apply on some samples (primary cells, *in vivo*…).

Chromatin Immunoprecipitation followed by Next Generation Sequencing (ChIP-Seq) is a technique of choice for studying protein-DNA interactions. ChIP has been used to pull down the Cas9 nuclease protein together with the DNA fragments to which the nuclease was bound (Kuscu et al., 2014; Wu et al., 2014). The immunoprecipitation of Cas9 bound to the genome is technically challenging due to the nuclease activity of Cas9. However, the introduction of two amino-acid changes (D10A and H840A) in Cas9-coding sequence results in a nuclease-inactive DNA-binding protein named “dead Cas9” (dCas9). Specific enrichment of dCas9 at on-target regions can be evaluated by ChIP-qPCR using ChIP-grade Cas9 antibodies (**Figure 4A**). Moreover, this approach can be extended to the unbiased analysis of off-target sites by ChIP-Seq (**Figure 4B**). dCas9-based ChIP-PCR/Seq is thus a powerful approach to score several sgRNA at once thanks to its rapidity, reduced sequencing cost and high coverage. Moreover, it is of predictive value of sgRNA performance upon association with catalytically active Cas9 (Kuscu et al., 2014), although Cas9 DNA-binding and cleavage activities are sometimes uncoupled (Wu et al., 2014). As no single method guarantees a complete coverage of off-target sites, multiple approaches should ideally be combined. Therefore, sequence-based *in silico* prediction combined to genome-wide ChIP-Seq dCas9-binding analysis can efficiently identify off-target sites (O’Geen et al., 2015).

**Figure 4.**
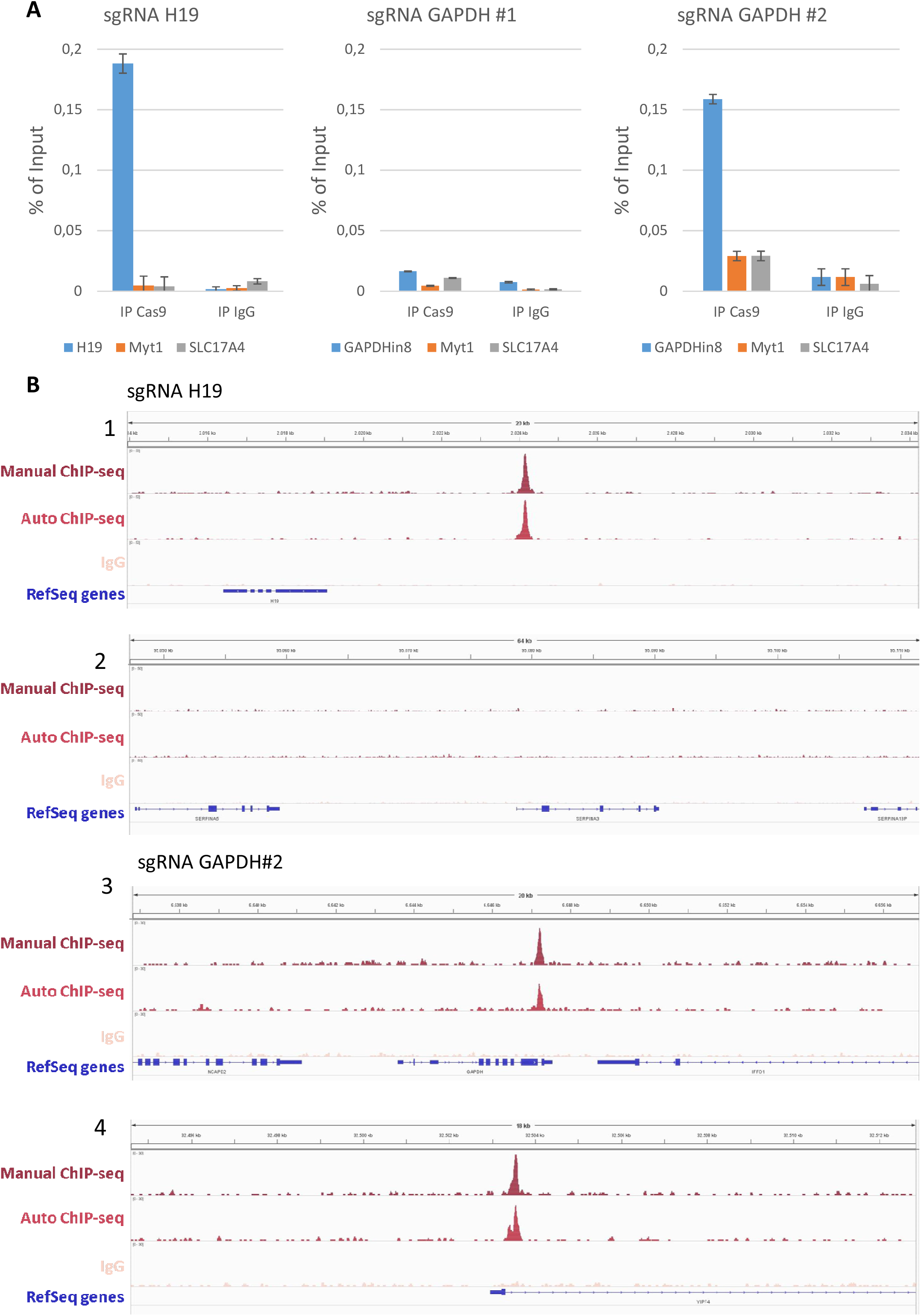
Evaluation of sgRNA specificity by dCas9 ChIP-PCR and ChIP-Seq. Sequences of sgRNA targeting the human H19 or GAPDH loci were designed using online selection tool (http://crispr.mit.edu/). The different sgRNAs (H19 sgRNA sequence: GTCTATCTCTGACAACCCTC; GAPDH sgRNA#1 sequence: GTCTGGCGCCCTCTGGTGGC and GAPDH sgRNA#2 sequence: AAAGACTCGGTCGGTGGTCT) were cloned in the pLenti-dCas9-2xAM plasmid (Addgene 92220, Fujita et al., 2018) and the resulting vectors were used to stably transfect HEK293T cells. ChIP was performed on sheared chromatin from 4 x 10 HEK293T cells using the iDeal ChIP-seq Kit for Transcription Factors (Diagenode Cat No. C01010170), 5μl of the polyclonal Cas9 antibody (Diagenode Cat No. C15310258) and 5μl of control IgG (Diagenode Cat No. C15410206). **A.** dCas9 ChIP-PCR analysis of one representative out of three independent experiments is shown (error bars show SD of replicates). Primers specific for the human H19, GAPDH, Myt1 and SLC17A4 were used for the qPCR. The figure shows the recovery, expressed as % of input (the relative amount of immunoprecipitated DNA compared to input DNA after qPCR analysis). **B.** dCas9 ChIP-Seq analysis. ChIP was performed manually or with the Auto iDeal ChIP-seq Kit for Transcription Factors (Diagenode, Cat. No. C01010172) and the IP-Star^®^ Compact Automated System (Diagenode Cat. No. B03000002) as described in the manual. Illumina compatible libraries were prepared on immunoprecipitated DNA using the MicroPlex Library Preparation Kit v2 according to the manual instructions (Diagenode Cat. No. C05010012). One nanogram of DNA was used per library preparation. The libraries were sequenced on HiSeq 3000 using SE 50pb reads. Cluster generation and sequencing were performed according to the manufacturer’s instructions. After trimming using Cutadapt, the reads were mapped to the human genome with the BWA aligner. Subsequent peak calling was performed using MACS. The figure shows the read distribution for the manual IP (top), the automated IP(middle) and IgG (bottom) samples. Peaks distribution of the three datasets in the region surrounding H19 (1) and a representative region of the genome (2) for HEK293T cells expressing a sgRNA for H19. Peaks distribution of the three datasets in the region surrounding GAPDH (3) and YIPF4 (4) for HEK293T cells expressing a sgRNA targeting GAPDH (sgRNA GAPDH #2).

Variants of dCas9 have recently been generated that allow repurposing of the system to a variety of applications. Fusing dCas9 to various transcriptional activating or repressing modules proved to be a potent way of regulating gene expression (Gilbert et al., 2013; Tanenbaum et al., 2014; Konermann et al., 2015; Yeo et al., 2018). This approach has also been used to identify enhancers of key loci (Simeonov et al., 2017). Moreover, dCas9 can be fused to domains that regulate the epigenetic landscape at endogenous loci (O’Geen et al., 2017). It can also be used to label endogenous loci for live visualization (Neguembor et al., 2017) or to edit a single base in the genome (Komor et al., 2016). In those applications, the binding specificity of dCas9 fused to various effectors could be tested by dCas9 ChIP-Seq as we describe here.

### Perspectives

From the first description of Cas9 derived from *Streptococcus Pyogenes* for gene editing in 2013, an incredible progress has been made to optimize and adapt its use in a wide range of applications. Structural studies of Cas9 led to the generation of several variants such as enhanced specificity Cas9 (eSpcas9), high fidelity Cas9 (Cas9-HF1) and hyper-accurate Cas9 (HypaCas9) which display increased specificity due to reduced DNA-binding affinity (eSpCas9 and Cas9-HF1) (Slaymaker et al., 2016; Kleinstiver et al., 2016) or locking of the nuclease domain upon guide/target mismatches (HypaCas9) (Chen et al., 2017). In addition, Cas9 nickases (Cas9n) were developed by inactivating the cleavage activity on target or non-target DNA and have been demonstrated to nick only one DNA strand instead of generating a double strand break (DSB). DSB are generated only upon recruitment of a Cas9n pair with two sgRNA that target opposite strands in close proximity (Hsu et al., 2013; Cho et al., 2014), thereby increasing specificity by double selection. A similar strategy was used to develop a fusion of dCas9 with the catalytic domain of FokI nuclease (fCas9) which induces DSB only upon dimerization of the FokI domains by sgRNA pairing to complementary strands (Tsai et al., 2014). With the same aim of reducing Cas9 off-target activity, several Cas9 variants whose editing activity can be irreversibly or reversibly programmed are now also available (Adli M, 2018; Wu et al., 2018 for review). Finally, engineered Cas9 variants with novel PAM specificities enlarge the edition spectrum to previously inaccessible sites (Kleinstiver et al., 2015 and 2016).

Several limitations have yet to be addressed to promote Cas9 use in gene therapy. First, the source of Cas9 nucleases, i.e. S. *pyogenes* and S. *aureus*, are common human pathogens. Recent reports have highlighted pre-existing immunity towards both SpCas9 and SaCas9 in the human population, with a high prevalence of both Cas9-reactive T cells and antibodies (Wagner et al., 2018; Simhadri et al., 2018; Charlesworth et al., 2018). Although it is still unclear whether AAV delivery of Cas9 leads to the immune rejection of transduced cells *in vivo*, strategies to control the anti-Cas9 T cell responses, such as transient immunosuppression or engineering Cas9 proteins with mutated T cell epitopes, are being considered (Crudele and Chamberlain, 2018; Ferdosi et al., 2018). Another limitation of Cas9 for its use in gene therapy resides in its rather large size which is incompatible for efficient packaging into Adeno-associated virus (AAV) vectors, the most commonly used delivery systems in gene therapy. Although this hurdle can be overcome by the separation of the recognition lobe from the nuclease lobe into two separate vectors (Truong et al., 2015), the emergence of Cas9 orthologs of smaller size might provide more efficient alternatives (Cebrian-Serrano et al., 2017, for review). Beside solving the delivery problem, CRISPR effectors from other bacterial and archeal species offer different substrate specificities or operate according to different mechanisms. This is notably the case of Cas12a (Cpf1) which is structurally different from Cas9, has no requirement for tracer RNA, recognizes a T-rich (TTTN) PAM sequence lying 5’ of the target sequence, and uses a different mechanism for target recognition and cleavage (Zetsche et al., 2015a). Cpf1 also possesses the ability to cleave RNA and generate multiple crRNAs from a single pre-crRNA array. This capacity has been harnessed to achieve multiplex gene editing using a single pre-crRNA array, which can both increase KO efficiency (when using multiple crRNAs targeting the same locus) or easily KO multiple genes with a single construct (Zetsche et al., 2017). Moreover, gene editing by Cpf1 results in lower off-target effects than Cas9, as evidenced by genome-wide analysis of edited cells (Kim et al., 2016). Finally, the discovery of Cas13a and CasRx as RNA-guided nucleases targeting RNA paves the way to new therapeutic approaches based on RNA editing (Shmakov et al., 2017, Konermann et al., 2018).

While a number of solutions and guidelines to harness CRISPR-Cas9 based gene targeting has been provided, it is expected that therapeutic, industrial, and research applications will still place high demand on improving the specificity and efficiency of the CRISPR/Cas9 system. As CRISPR-based gene targeting technology continues to become more sophisticated and diverse, optimized procedures and quality controls guidelines should be established.

## Acknowledgements

We thank Romuald Soin and Nadège Delacourt for technical support. Claude Van Campenhout was funded by a FIRST Entreprise grant from DGO6 from the Walloon Region, Belgium.

## References

Adli M. The CRISPR tool kit for genome editing and beyond. Nat Commun. 2018 May 15;9(1):1911. doi: 10.1038/s41467-018-04252-2

Anders C, Jinek M. In vitro Enzymology of Cas9. Methods Enzymol. 2014; 546:1–20.

Ayadi A, Birling MC, Bottomley J, Bussell J, Fuchs H, Fray M, Gailus-Durner V, Greenaway S, Houghton R, Karp N, Leblanc S, Lengger C, Maier H, Mallon AM, Marschall S, Melvin D, Morgan H, Pavlovic G, Ryder E, Skarnes WC, Selloum M, Ramirez-Solis R, Sorg T, Teboul L, Vasseur L, Walling A, Weaver T, Wells S, White JK, Bradley A, Adams DJ, Steel KP, Hrabě de Angelis M, Brown SD, Herault Y. Mouse large-scale phenotyping initiatives: overview of the European Mouse Disease Clinic (EUMODIC) and of the Wellcome Trust Sanger Institute Mouse Genetics Project. Mamm Genome. 2012 Oct;23(9–10):600–10.

Barrangou, R., Fremaux, C., Deveau, H., Richards, M., Boyaval, P., Moineau, S., Romero, D.A., and Horvath, P. CRISPR provides acquired resistance against viruses in prokaryotes. Science 2007; 315:1709–1712.

Burger A, Lindsay H, Felker A, Hess C, Anders C, Chiavacci E, Zaugg J, Weber LM, Catena R, Jinek M, Robinson MD, Mosimann C. Maximizing mutagenesis with solubilized CRISPR-Cas9 ribonucleoprotein complexes. Development. 2016 Jun 1;143(11):2025–37.

Cameron P, Fuller CK, Donohoue PD, Jones BN, Thompson MS, Carter MM, Gradia S, Vidal B, Garner E, Slorach EM, Lau E, Banh LM, Lied AM, Edwards LS, Settle AH, Capurso D, Llaca V, Deschamps S, Cigan M, Young JK, May AP. Mapping the genomic landscape of CRISPR-Cas9 cleavage. Nat Methods. 2017 Jun;14(6):600–606.

Cao J, Wu L, Zhang SM, Lu M, Cheung WK, Cai W, Gale M, Xu Q, Yan Q. An easy and efficient inducible CRISPR/Cas9 platform with improved specificity for multiple genetargeting. Nucleic Acids Res. 2016 Nov 2;44(19):e149.

Cebrian-Serrano A, Davies B. CRISPR-Cas orthologues and variants: optimizing the repertoire, specificity and delivery of genome engineering tools. Mamm Genome. 2017 Aug;28(7–8):247–261.

Challa AK, Boitet ER, Turner AN, Johnson LW, Kennedy D, Downs ER, Hymel KM, Gross AK, Kesterson RA. Novel Hypomorphic Alleles of the Mouse Tyrosinase Gene Induced by CRISPR-Cas9 Nucleases Cause Non-Albino Pigmentation Phenotypes. PLoS One 2016 May 25;11 (5):e0155812.

Charlesworth CT, Deshpande PS, Dever DP, Dejene B, Gomez-Ospina N, Mantri S, Pavel-Dinu M, Camarena J, Weinberg KI, Porteus MH. Identification of Pre-Existing Adaptive Immunity to Cas9 Proteins in Humans. BioRxiv 2018 doi: https://doi.org/10.1101/243345

Chen JS, Dagdas YS, Kleinstiver BP, Welch MM, Sousa AA, Harrington LB, Sternberg SH, Joung JK, Yildiz A, Doudna JA. Enhanced proofreading governs CRISPR-Cas9 targeting accuracy. Nature 2017 Oct 19;550(7676):407–410.

Chen X, Liu J, Janssen JM, Gonçalves MAFV. The Chromatin Structure Differentially Impacts High-Specificity CRISPR-Cas9 Nuclease Strategies. Mol Ther Nucleic Acids 2017 Sep 15;8:558–563.

Chen X, Rinsma M, Janssen JM, Liu J, Maggio I, Gonçalves MA. Probing the impact of chromatin conformation on genome editing tools. Nucleic Acids Res. 2016 Jul 27;44(13):6482–92.

Chen Y, Zeng S, Hu R, Wang X, Huang W, Liu J, Wang L, Liu G, Cao Y, Zhang Y. Using local chromatin structure to improve CRISPR/Cas9 efficiency in zebrafish. PLoS One 2017 Aug 11;12(8):e0182528.

Cho SW, Kim S, Kim JM, Kim JS. Targeted genome engineering in human cells with the Cas9 RNA-guided endonuclease. Nat Biotechnol. 2013 Mar;31(3):230–2.

Cho SW, Kim S, Kim Y, Kweon J, Kim HS, Bae S, Kim JS. Analysis of off-target effects of CRISPR/Cas-derived RNA-guided endonucleases and nickases. Genome Res. 2014 Jan;24(1):132–41.

Cong, L., Ran, F.A., Cox, D., Lin, S., Barretto, R., Habib, N., Hsu, P.D., Wu, X., Jiang, W., Marraffini, L.A., et al. Multiplex genome engineering using CRISPR/Cas systems. Science 2013 339, 819–823.

Courtney DG, Moore JE, Atkinson SD, Maurizi E, Allen EH, Pedrioli DM, McLean WH, Nesbit MA, Moore CB. CRISPR/Cas9 DNA cleavage at SNP-derived PAM enables both in vitro and in vivo KRT12 mutation-specific targeting. Gene Ther. 2016 Jan;23(1):108–12.

Crudele JM, Chamberlain JS. Cas9 immunity creates challenges for CRISPR gene editing therapies. Nat Commun. 2018 Aug 29;9(1):3497. doi: 10.1038/s41467-018-05843-9.

Daer RM, Cutts JP, Brafman DA, Haynes KA. The Impact of Chromatin Dynamics on Cas9-Mediated Genome Editing in Human Cells. ACS Synth Biol. 2017 Mar 17;6(3):428–438.

D’Astolfo DS, Pagliero RJ, Pras A, Karthaus WR, Clevers H, Prasad V, Lebbink RJ, Rehmann H, Geijsen N. Efficient intracellular delivery of native proteins. Cell 2015 Apr 23;161(3):674–690.

Deltcheva E, Chylinski K, Sharma CM, Gonzales K, Chao Y, Pirzada ZA, Eckert MR, Vogel J, Charpentier E. CRISPR RNA maturation by trans-encoded small RNA and host factor RNase III. Nature 2011 Mar 31;471(7340):602–7.

Driever W, Solnica-Krezel L, Schier AF, Neuhauss SC, Malicki J, Stemple DL, Stainier DY, Zwartkruis F, Abdelilah S, Rangini Z, Belak J, Boggs C. A genetic screen for mutations affecting embryogenesis in zebrafish. Development 1996 Dec;123:37–46.

Doench JG, Fusi N, Sullender M, Hegde M, Vaimberg EW, Donovan KF, Smith I, Tothova Z^3^, Wilen C, Orchard R, Virgin HW, Listgarten J, Root DE. Optimized sgRNA design to maximize activity and minimize off-target effects of CRISPR-Cas9. Nat Biotechnol. 2016 Feb;34(2):184–191. doi: 10.1038/nbt.3437.

El-Brolosy MA and Stainier DYR. Genetic compensation: A phenomenon in search of mechanisms. PloS Genet. 2017 Jul 13;13(7):e1006780.

Endo M, Mikami M, Toki S. Multigene knockout utilizing off-target mutations of the CRISPR/Cas9 system in rice. Plant Cell Physiol. 2015 Jan;56(1):41–7.

Eubelen M, Bostaille N, Cabochette P, Gauquier A, Tebabi P, Dumitru AC, Koehler M, Gut P, Alsteens D, Stainier DYR, Garcia-Pino A, Vanhollebeke B. A molecular mechanism for Wnt ligand-specific signaling. Science 2018 Aug 17; 361: (6403) pii: eaat1178.

Ferdosi SR, Ewaisha R, Moghadam F, Krishna S, Park JG, Ebrahimkhani MR, Kiani S, Anderson KS. Multifunctional CRISPR/Cas9 with engineered immunosilenced human T cell epitopes. BioRxiv 2018 doi: https://doi.org/10.1101/360198

Frock RL, Hu J, Meyers RM, Ho YJ, Kii E, Alt FW. Genome-wide detection of DNA double-stranded breaks induced by engineered nucleases. Nat Biotechnol. 2015 Feb;33(2):179–86.

Fu Y, Foden JA, Khayter C, Maeder ML, Reyon D, Joung JK, Sander JD. High-frequency off-target mutagenesis induced by CRISPR-Cas nucleases in human cells. Nat Biotechnol. 2013 Sep;31(9):822–6.

Fu Y, Sander JD, Reyon D, Cascio VM, Joung JK. Improving CRISPR-Cas nuclease specificity using truncated guide RNAs. Nat Biotechnol. 2014 Mar;32(3):279–284.

Fujita T, Yuno M, Fujii H. enChIP systems using different CRISPR orthologues and epitope tags. BMC Res Notes 2018 Feb 27;11 (1):154.

Gagnon JA, Valen E, Thyme SB, Huang P, Akhmetova L, Pauli A, Montague TG, Zimmerman S, Richter C, Schier AF. Efficient mutagenesis by Cas9 protein-mediated oligonucleotide insertion and large-scale assessment of single-guide RNAs. PLoS One 2014 May 29;9(5):e98186.

Gasiunas G, Barrangou R, Horvath P, Siksnys V. Cas9-crRNA ribonucleoprotein complex mediates specific DNA cleavage for adaptive immunity in bacteria. Proc Natl Acad Sci U S A. 2012 Sep 25;109(39):E2579–86.

Gilbert LA, Larson MH, Morsut L, Liu Z, Brar GA, Torres SE, Stern-Ginossar N, Brandman O, Whitehead EH, Doudna JA, Lim WA, Weissman JS, Qi LS. CRISPR-mediated modular RNA-guided regulation of transcription in eukaryotes. Cell 2013 Jul 18;154(2):442–51.

Goto T, Hara H., Nakauchi H., Hochi S., Hirabayashi M. Hypomorphic phenotype of Foxn1 gene-modified rats by CRISPR/Cas9 system. Transgenic Res. 2016 25, 533–544.

Haeussler M, Schönig K, Eckert H, Eschstruth A, Mianné J, Renaud JB, Schneider-Maunoury S, Shkumatava A, Teboul L, Kent J, Joly JS, Concordet JP. Evaluation of off-target and on-target scoring algorithms and integration into the guide RNA selection tool CRISPOR. Genome Biol. 2016 Jul 5;17(1): 148.

Haffter P, Granato M, Brand M, Mullins MC, Hammerschmidt M, Kane DA, Odenthal J, van Eeden FJ, Jiang YJ, Heisenberg CP, Kelsh RN, Furutani-Seiki M, Vogelsang E, Beuchle D, Schach U, Fabian C, Nüsslein-Volhard C. The identification of genes with unique and essential functions in the development of the zebrafish, Danio rerio. Development 1996 Dec;123:1–36.

Hamada H, Liu Y, Nagira Y, Miki R, Taoka N, Imai R. Biolistic-delivery-based transient CRISPR/Cas9 expression enables in planta genome editing in wheat. Sci Rep. 2018 Sep 26;8(1):14422.

Hart T, Chandrashekhar M, Aregger M, Steinhart Z, Brown KR, MacLeod G, Mis M, Zimmermann M, Fradet-Turcotte A, Sun S, Mero P, Dirks P, Sidhu S, Roth FP, Rissland OS, Durocher D, Angers S, Moffat J. High-Resolution CRISPR Screens Reveal Fitness Genes and Genotype-Specific Cancer Liabilities. Cell 2015 Dec 3;163(6): 1515–26.

He Z, Shi X, Liu M, Sun G, Proudfoot C, Whitelaw CB, Lillico SG, Chen Y. Comparison of surrogate reporter systems for enrichment of cells with mutations induced by genome editors. J Biotechnol. 2016 Mar 10;221:49–54.

Horlbeck MA, Witkowsky LB, Guglielmi B, Replogle JM, Gilbert LA, Villalta JE, Torigoe SE, Tjian R, Weissman JS. Nucleosomes impede Cas9 access to DNA in vivo and in vitro. Elife 2016 Mar 17;5. pii: e12677.

Hsu PD, Scott DA, Weinstein JA, Ran FA, Konermann S, Agarwala V, Li Y, Fine EJ, Wu X, Shalem O, Cradick TJ, Marraffini LA, Bao G, Zhang F. DNA targeting specificity of RNA-guided Cas9 nucleases. Nat Biotechnol. 2013 Sep;31(9):827–32.

Hyams G, Abadi S, Lahav S, Avni A, Halperin E, Shani E, Mayrose I. CRISPys: Optimal sgRNA Design for Editing Multiple Members of a Gene Family Using the CRISPR System. J Mol Biol. 2018 Jul 20;430(15):2184–2195. doi: 10.1016/j.jmb.2018.03.019.

Isaac RS, Jiang F, Doudna JA, Lim WA, Narlikar GJ, Almeida R. Nucleosome breathing and remodeling constrain CRISPR-Cas9 function. Elife 2016 Apr 28;5. pii: e13450.

Jao LE, Wente SR, Chen W. Efficient multiplex biallelic zebrafish genome editing using a CRISPR nuclease system. Proc Natl Acad Sci U S A. 2013 Aug 20;110(34):13904–9.

Jensen KT, Fløe L, Petersen TS, Huang J, Xu F, Bolund L, Luo Y, Lin L. Chromatin accessibility and guide sequence secondary structure affect CRISPR-Cas9 gene editing efficiency. FEBS Lett. 2017 Jul;591 (13):1892–1901.

Jinek, M., Chylinski, K., Fonfara, I., Hauer, M., Doudna, J.A., and Charpentier, E. (2012). A programmable dual-RNA-guided DNA endonuclease in adaptive bacterial immunity. Science 337, 816–821.

Kim D, Bae S, Park J, Kim E, Kim S, Yu HR, Hwang J, Kim JI, Kim JS. Digenome-seq: genome-wide profiling of CRISPR-Cas9 off-target effects in human cells. Nat Methods 2015 Mar;12(3):237–43.

Kim D, Kim J, Hur JK, Been KW, Yoon SH, Kim JS. Genome-wide analysis reveals specificities of Cpf1 endonucleases in human cells. Nat Biotechnol. 2016 Aug;34(8):863–8.

Kim K, Park SW, Kim JH, Lee SH, Kim D, Koo T, Kim KE, Kim JH, Kim JS. Genome surgery using Cas9 ribonucleoproteins for the treatment of age-related macular degeneration. Genome Res. 2017 Mar;27(3):419–426.

Kim S, Kim D, Cho SW, Kim J, Kim JS. Highly efficient RNA-guided genome editing in human cells via delivery of purified Cas9 ribonucleoproteins. Genome Res. 2014 Jun;24(6):1012–9.

Kleinstiver BP, Prew MS, Tsai SQ, Topkar VV, Nguyen NT, Zheng Z, Gonzales AP, Li Z, Peterson RT, Yeh JR, Aryee MJ, Joung JK. Engineered CRISPR-Cas9 nucleases with altered PAM specificities. Nature 2015 Jul 23;523(7561):481–5.

Kleinstiver BP, Pattanayak V, Prew MS, Tsai SQ, Nguyen NT, Zheng Z, Joung JK. High-fidelity CRISPR-Cas9 nucleases with no detectable genome-wide off-target effects. Nature 2016 Jan 28;529(7587):490–5.

Komor AC, Kim YB, Packer MS, Zuris JA, Liu DR. Programmable editing of a target base in genomic DNA without double-stranded DNA cleavage. Nature 2016 May 19;533(7603):420–4.

Konermann S, Brigham MD, Trevino AE, Joung J, Abudayyeh OO, Barcena C, Hsu PD, Habib N, Gootenberg JS, Nishimasu H, Nureki O, Zhang F. Genome-scale transcriptional activation by an engineered CRISPR-Cas9 complex. Nature. 2015 Jan 29;517(7536):583–8.

Konermann S, Lotfy P, Brideau NJ, Oki J, Shokhirev MN, Hsu PD. Transcriptome Engineering with RNA-Targeting Type VI-D CRISPR Effectors. Cell 2018 Apr 19;173(3):665–676.e14.

Kuscu C, Arslan S, Singh R, Thorpe J, Adli M. Genome-wide analysis reveals characteristics of off-target sites bound by the Cas9 endonuclease. Nat Biotechnol. 2014 Jul;32(7):677–83.

Landry JJ, Pyl PT, Rausch T, Zichner T, Tekkedil MM, Stütz AM, Jauch A, Aiyar RS, Pau G, Delhomme N, Gagneur J, Korbel JO, Huber W, Steinmetz LM. The genomic and transcriptomic landscape of a HeLa cell line. G3 (Bethesda) 2013 Aug 7;3(8):1213–24.

Lenoir WF, Lim TL, Hart T. PICKLES: the database of pooled in-vitro CRISPR knockout library essentiality screens. Nucleic Acids Res. 2018 Jan 4;46(D1):D776–D780.

Li Y, Mendiratta S, Ehrhardt K, Kashyap N, White MA, Bleris L. Exploiting the CRISPR/Cas9 PAM Constraint for Single-Nucleotide Resolution Interventions. PLoS One 2016 Jan 20;11(1):e0144970.

Liang X, Potter J, Kumar S, Zou Y, Quintanilla R, Sridharan M, Carte J, Chen W, Roark N, Ranganathan S, Ravinder N, Chesnut JD. Rapid and highly efficient mammalian cell engineering via Cas9 protein transfection. J Biotechnol. 2015 Aug 20;208:44–53.

Mali P, Yang L, Esvelt KM, Aach J, Guell M, DiCarlo JE, Norville JE, Church GM. RNA-guided human genome engineering via Cas9. Science 2013 Feb 15;339(6121):823–6.

Mircetic J, Steinebrunner I, Ding L, Fei JF, Bogdanova A, Drechsel D, Buchholz F. Purified Cas9 Fusion Proteins for Advanced Genome Manipulation. Advanced Science News Small Methods 2017 1; 1600052

Mojica, F.J.M., Diez-Villasenor, C.S., Garcia-Martinez, J.S., and Soria, E. Intervening Sequences of Regularly Spaced Prokaryotic Repeats Derive from Foreign Genetic Elements. J Mol Evol. 2005 60, 174–182.

Neguembor MV, Sebastian-Perez R, Aulicino F, Gomez-Garcia PA, Cosma MP, Lakadamyali M. (Po)STAC (Polycistronic SunTAg modified CRISPR) enables live-cell and fixed-cell super-resolution imaging of multiple genes. Nucleic Acids Res. 2018 46 (5) e30.

O’Geen H, Henry IM, Bhakta MS, Meckler JF, Segal DJ. A genome-wide analysis of Cas9 binding specificity using ChIP-seq and targeted sequence capture. Nucleic Acids Res. 2015 Mar 31;43(6):3389–404.

O’Geen H, Ren C, Nicolet CM, Perez AA, Halmaj J, Le VM, Mackay JP, Farnham PJ, Segal DJ. dCas9-based epigenome editing suggests acquisition of histone methylation is not sufficient for target gene repression. Nucleic Acids Res. 2017 Sep 29;45(17):9901–9916.

Qi LS, Larson MH, Gilbert LA, Doudna JA, Weissman JS, Arkin AP, Lim WA. Repurposing CRISPR as an RNA-guided platform for sequence-specific control of gene expression. Cell 2013 Feb 28;152(5):1173–83.

Ramakrishna S, Kwaku Dad AB, Beloor J, Gopalappa R, Lee SK, Kim H. Gene disruption by cell-penetrating peptide-mediated delivery of Cas9 protein and guide RNA. Genome Res. 2014a Jun;24(6):1020–7.

Ramakrishna S, Cho SW, Kim S, Song M, Gopalappa R, Kim JS, Kim H. Surrogate reporter-based enrichment of cells containing RNA-guided Cas9 nuclease-induced mutations. Nat Commun. 2014b Feb 26;5:3378.

Ran FA, Cong L, Yan WX, Scott DA, Gootenberg JS, Kriz AJ, Zetsche B, Shalem O, Wu X, Makarova KS, Koonin EV, Sharp PA, Zhang F. In vivo genome editing using Staphylococcus aureus Cas9. Nature 2015 Apr 9;520(7546):186–91.

Sentmanat MF, Peters ST, Florian CP, Connelly JP, Pruett-Miller SM. A Survey of Validation Strategies for CRISPR-Cas9 Editing. Sci Rep. 2018 Jan 17;8(1):888.

Shi J, Wang E, Milazzo JP, Wang Z, Kinney JB & Vakoc CR. Discovery of cancer drug targets by CRISPR-Cas9 screening of protein domains. Nat Biotechnol. 2015 33, 661–667.

Shifrut E, Carnevale J, Tobin V, Roth TL, Woo JM, Bui CT, Li PJ, Diolaiti ME, Ashworth A, Marson A. Genome-wide CRISPR Screens in Primary Human T Cells Reveal Key Regulators of Immune Function. Cell. 2018 Nov 13. pii: S0092–8674(18)31333–3.

Shmakov S, Smargon A, Scott D, Cox D, Pyzocha N, Yan W, Abudayyeh OO, Gootenberg JS, Makarova KS, Wolf YI, Severinov K, Zhang F, Koonin EV. Diversity and evolution of class 2 CRISPR-Cas systems. Nat Rev Microbiol. 2017 Mar;15(3):169–182.

Simeonov DR, Gowen BG, Boontanrart M, Roth TL, Gagnon JD, Mumbach MR, Satpathy AT, Lee Y, Bray NL, Chan AY, Lituiev DS, Nguyen ML, Gate RE, Subramaniam M, Li Z, Woo JM, Mitros T, Ray GJ, Curie GL, Naddaf N, Chu JS, Ma H, Boyer E, Van Gool F, Huang H, Liu R, Tobin VR, Schumann K, Daly MJ, Farh KK, Ansel KM, Ye CJ, Greenleaf WJ, Anderson MS, Bluestone JA, Chang HY, Corn JE, Marson A. Discovery of stimulation-responsive immune enhancers with CRISPR activation. Nature. 2017 Sep 7;549(7670):111–115.

Simhadri VL, McGill J, McMahon S, Wang J, Jiang H, Sauna ZE. Prevalence of Pre-existing Antibodies to CRISPR-Associated Nuclease Cas9 in the USA Population. Mol Ther Methods Clin Dev. 2018 Jun 15;10:105–112.

Slaymaker IM, Gao L, Zetsche B, Scott DA, Yan WX, Zhang F. Rationally engineered Cas9 nucleases with improved specificity. Science 2016 Jan 1;351(6268):84–8.

Smith C, Gore A, Yan W, Abalde-Atristain L, Li Z, He C, Wang Y, Brodsky RA, Zhang K, Cheng L, Ye Z. Whole-genome sequencing analysis reveals high specificity of CRISPR/Cas9 and TALEN-based genome editing in human iPSCs. Cell Stem Cell. 2014 Jul 3;15(1):12–3.

Song J, Yang D, Ruan J, Zhang J, Chen YE, Xu J. Production of immunodeficient rabbits by multiplex embryo transfer and multiplex gene targeting. Sci Rep. 2017 Sep 22;7(1):12202.

Svitashev S, Schwartz C, Lenderts B, Young JK, Mark Cigan A. Genome editing in maize directed by CRISPR-Cas9 ribonucleoprotein complexes. Nat Commun. 2016 Nov 16;7:13274.

Thyme SB, Akhmetova L, Montague TG, Valen E, Schier AF. Internal guide RNA interactions interfere with Cas9-mediated cleavage. Nat Commun. 2016 Jun 10;7:11750.

Ting PY, Parker AE, Lee JS, Trussell C, Sharif O, Luna F, Federe G, Barnes SW, Walker JR, Vance J, Gao MY, Klock HE, Clarkson S, Russ C, Miraglia LJ, Cooke MP, Boitano AE, McNamara P, Lamb J, Schmedt C, Snead JL. Guide Swap enables genome-scale pooled CRISPR-Cas9 screening in human primary cells. Nat Methods. 2018 Nov;15(11):941–946.

Truong DJ, Kühner K, Kühn R, Werfel S, Engelhardt S, Wurst W, Ortiz O. Development of an intermediated split-Cas9 system for gene therapy. Nucleic Acids Res. 2015 Jul 27;43(13):6450–8.

Tsai SQ, Wyvekens N, Khayter C, Foden JA, Thapar V, Reyon D, Goodwin MJ, Aryee MJ, Joung JK. Dimeric CRISPR RNA-guided FokI nucleases for highly specific genome editing. Nat Biotechnol. 2014 Jun;32(6):569–76.

Tsai SQ, Zheng Z, Nguyen NT, Liebers M, Topkar VV, Thapar V, Wyvekens N, Khayter C, Iafrate AJ, Le LP, Aryee MJ, Joung JK. GUIDE-seq enables genome-wide profiling of off-target cleavage by CRISPR-Cas nucleases. Nat Biotechnol. 2015 Feb;33(2):187–197.

Tsai SQ, Nguyen NT, Malagon-Lopez J, Topkar VV, Aryee MJ, Joung JK. CIRCLE-seq: a highly sensitive in vitro screen for genome-wide CRISPR-Cas9 nuclease off-targets. Nat Methods 2017 Jun;14(6):607–614.

Ulrich F, Carretero-Ortega J, Menéndez J, Narvaez C, Sun B, Lancaster E, Pershad V, Trzaska S, Véliz E, Kamei M, Prendergast A, Kidd KR, Shaw KM, Castranova DA, Pham VN, Lo BD, Martin BL, Raible DW, Weinstein BM, Torres-Vazquez J. Reck enables cerebrovascular development by promoting canonical Wnt signalling. Development 2016 Jan 1; 143(1):147–59.

Uusi-Mäkelä MIE, Barker HR, Bäuerlein CA, Häkkinen T, Nykter M, Rämet M. Chromatin accessibility is associated with CRISPR-Cas9 efficiency in the zebrafish (Danio rerio). PLoS One 2018 Apr 23;13(4):e0196238.

Vanhollebeke B, Stone OA, Bostaille N, Cho C, Zhou Y, Maquet E, Gauquier A, Cabochette P, Fukuhara S, Mochizuki N, Nathans J, Stainier DY. Tip cell-specific requirement for an atypical Gpr124- and Reck-dependent Wnt/β-catenin pathway during brain angiogenesis. Elife 2015 Jun 8;4.

Vouillot L, Thélie A, Pollet N. Comparison of T7E1 and surveyor mismatch cleavage assays to detect mutations triggered by engineered nucleases. G3 (Bethesda) 2015 Jan 7;5(3):407–15.

Wagner DL, Amini L, Wendering DJ, Burkhardt LM, Akyüz L, Reinke P, Volk HD, Schmueck-Henneresse M. High prevalence of Streptococcus pyogenes Cas9-reactive T cells within the adult human population. Nat Med. 2018 Oct 29. doi: 10.1038/s41591-018-0204-6.

Wang T, Birsoy K, Hughes NW, Krupczak KM, Post Y, Wei JJ, Lander ES, Sabatini DM. Identification and characterization of essential genes in the human genome. Science 2015 Nov 27;350(6264):1096–101.

Wang T, Wei JJ, Sabatini DM, Lander ES. Genetic screens in human cells using the CRISPR-Cas9 system. Science. 2014 Jan 3;343(6166):80–4. doi: 10.1126/science.1246981.

Wang X, Wang Y, Wu X, Wang J, Wang Y, Qiu Z, Chang T, Huang H, Lin RJ, Yee JK. Unbiased detection of off-target cleavage by CRISPR-Cas9 and TALENs using integrase-defective lentiviral vectors. Nat Biotechnol. 2015 Feb;33(2):175–8.

Wu X, Scott DA, Kriz AJ, Chiu AC, Hsu PD, Dadon DB, Cheng AW, Trevino AE, Konermann S, Chen S, Jaenisch R, Zhang F, Sharp PA. Genome-wide binding of the CRISPR endonuclease Cas9 in mammalian cells. Nat Biotechnol. 2014 Jul;32(7):670–6.

Wroblewska A, Dhainaut M, Ben-Zvi B, Rose SA, Park ES, Amir ED, Bektesevic A, Baccarini A, Merad M, Rahman AH, Brown BD. Protein Barcodes Enable High-Dimensional Single-Cell CRISPR Screens. Cell 2018 Oct 16. pii: S0092–8674(18)31234–0.

Wu WY, Lebbink JHG, Kanaar R, Geijsen N, van der Oost J. Genome editing by natural and engineered CRISPR-associated nucleases. Nat Chem Biol. 2018 Jul;14(7):642–651.

Wu Y, Xu K, Ren C, Li X, Lv H, Han F, Wei Z, Wang X, Zhang Z. Enhanced CRISPR/Cas9-mediated biallelic genome targeting with dual surrogate reporter-integrated donors. FEBS Lett. 2017 Mar;591(6):903–913.

Yeo NC, Chavez A, Lance-Byrne A, Chan Y, Menn D, Milanova D, Kuo CC, Guo X, Sharma S, Tung A, Cecchi RJ, Tuttle M, Pradhan S, Lim ET, Davidsohn N, Ebrahimkhani MR, Collins JJ, Lewis NE, Kiani S, Church GM. An enhanced CRISPR repressor for targeted mammalian gene regulation. Nat Methods 2018 Aug;15(8):611–616.

Zetsche B, Gootenberg JS, Abudayyeh OO, Slaymaker IM, Makarova KS, Essletzbichler P, Volz SE, Joung J, van der Oost J, Regev A, Koonin EV, Zhang F. Cpf1 is a single RNA-guided endonuclease of a class 2 CRISPR-Cas system. Cell. 2015a Oct 22;163(3):759–71.

Zetsche B, Volz SE, Zhang F. A split-Cas9 architecture for inducible genome editing and transcription modulation. Nat. Biotechnol. 2015b 33:139–142.

Zheng T, Hou Y, Zhang P, Zhang Z, Xu Y, Zhang L, Niu L, Yang Y, Liang D, Yi F, Peng W, Feng W, Yang Y, Chen J, Zhu YY, Zhang LH, Du Q. Profiling single-guide RNA specificity reveals a mismatch sensitive core sequence. Sci Rep. 2017 Jan 18;7:40638.

Zuris JA, Thompson DB, Shu Y, Guilinger JP, Bessen JL, Hu JH, Maeder ML, Joung JK, Chen ZY, Liu DR. Cationic lipid-mediated delivery of proteins enables efficient protein-based genome editing *in vitro* and *in vivo*. Nat Biotechnol. 2015 Jan;33(1):73–80.

